# Multimodal state-dependent connectivity analysis of arousal and autonomic centers in the brainstem and basal forebrain

**DOI:** 10.1101/2024.11.11.623092

**Authors:** Haatef Pourmotabbed, Caroline G. Martin, Sarah E. Goodale, Derek J. Doss, Shiyu Wang, Roza G. Bayrak, Hakmook Kang, Victoria L. Morgan, Dario J. Englot, Catie Chang

## Abstract

Vigilance is a continuously altering state of cortical activation that influences cognition and behavior and is disrupted in multiple brain pathologies. Neuromodulatory nuclei in the brainstem and basal forebrain are implicated in arousal regulation and are key drivers of widespread neuronal activity and communication. However, it is unclear how their large-scale brain network architecture changes across dynamic variations in vigilance state (i.e., alertness and drowsiness). In this study, we leverage simultaneous EEG and 3T multi-echo functional magnetic resonance imaging (fMRI) to elucidate the vigilance-dependent connectivity of arousal regulation centers in the brainstem and basal forebrain. During states of low vigilance, most of the neuromodulatory nuclei investigated here exhibit a stronger global correlation pattern and greater connectivity to the thalamus, precuneus, and sensory and motor cortices. In a more alert state, the nuclei exhibit the strongest connectivity to the salience, default mode, and auditory networks. These vigilance-dependent correlation patterns persist even after applying multiple preprocessing strategies to reduce systemic vascular effects. To validate our findings, we analyze two large 3T and 7T fMRI datasets from the Human Connectome Project and demonstrate that the static and vigilance-dependent connectivity profiles of the arousal nuclei are reproducible across 3T multi-echo, 3T single-echo, and 7T single-echo fMRI modalities. Overall, this work provides novel insights into the role of neuromodulatory systems in vigilance-related brain activity.

## 1. INTRODUCTION

Vigilance is a continuously altering state of physiological and psychological activation (i.e., alertness and drowsiness) that impacts the ability of the brain to process information and respond to external stimuli (Oken et al., 2006; Sara and Bouret, 2012). Higher levels of alertness result in enhanced cognitive processing, greater emotional reactivity, and an improved capability for sustained attention (Canales-Johnson et al., 2020; Franzen et al., 2008; Jagannathan et al., 2022). Additionally, impairments of vigilance occur in multiple brain pathologies and contribute to the development of neurocognitive deficits in executive function and attention. These vigilance impairments include hyperarousal in neuropsychiatric disorders (Hegerl and Hensch, 2014; Xie et al., 2024) and excessive daytime sleepiness and sleep-wake disturbances in traumatic brain injury, epilepsy, Alzheimer’s disease, and Parkinson’s disease (Englot et al., 2020; Rothman and Mattson, 2012; Sandsmark et al., 2017). Identifying the neural circuit mechanisms underlying alterations in vigilance state may aid in uncovering novel therapies for neurocognitive deficits in various brain disorders.

Key drivers of widespread neuronal activity and communication include neuromodulatory centers in the brainstem and basal forebrain (van den Brink et al., 2019). These neuromodulatory nuclei consist of monoaminergic, glutamatergic, and cholinergic neurons that project to the thalamus, hypothalamus, and widespread areas of the cortex, mediating cortical activation and autonomic function (Brown et al., 2012; Edlow et al., 2012; Scammell et al., 2017; Zaborszky et al., 2008). Human and animal studies have provided evidence that the neuronal activity of neuromodulatory nuclei is associated with changes in widespread cortical activity, brain network organization, and markers of arousal and attention (Grimm et al., 2024; Liu et al., 2018; Taylor et al., 2022; Zerbi et al., 2019). For instance, blood oxygenation level dependent (BOLD) signals in the locus coeruleus (LC) and nucleus basalis of Meynert (NBM) have been shown to be correlated with pupil diameter, low-frequency electrophysiological activity, and attentional task response (Joshi et al., 2016; Liu et al., 2018; Murphy et al., 2014). Furthermore, pharmacological studies have found that inactivation of neurons in the NBM leads to suppression of global brain signals (Turchi et al., 2018) and modulation of monoamine neurotransmitters results in altered resting-state functional connectivity (FC) (van den Brink et al., 2016).

Neuroimaging studies in healthy individuals have sought to characterize the structural and functional connectivity of neuromodulatory nuclei in the brainstem and basal forebrain (Bar et al., 2016; Beliveau et al., 2015; Cauzzo et al., 2022; Hansen et al., 2024; Yuan et al., 2019; Zhang et al., 2016). Abnormalities in the connectivity of these subcortical regions have also been observed in multiple neurological conditions, suggesting that mapping of the FC may provide a valuable avenue for identifying brain targets for therapeutic neuromodulation (Edlow et al., 2021; Englot et al., 2020; Gonzalez et al., 2021; Kelberman et al., 2020; Serra et al., 2018). However, to date, functional magnetic resonance imaging (fMRI) studies have not comprehensively mapped vigilance-related alterations in the FC of the brainstem and basal forebrain. Dynamic changes in the spatiotemporal activity and FC of the cortex have been linked to altering states of alertness and wakefulness (Liu and Falahpour, 2020; Martin et al., 2021).

These state-dependent effects are often unaccounted for due to the difficulty in estimating vigilance based on fMRI alone (Goodale et al., 2021; Liu and Falahpour, 2020; Martin et al., 2021). Subcortical neuromodulatory systems may be involved in coordinating arousal changes in the cortex (Brown et al., 2012; Scammell et al., 2017), and characterizing the vigilance-dependent connectivity of the subcortical activating structures can provide novel insights into their role in regulating brain activity. Therefore, in this study, we leveraged simultaneously recorded electroencephalography (EEG) and fMRI data to elucidate the functional network architecture of neuromodulatory nuclei in different vigilance states. The EEG data were used to identify time periods of alertness and drowsiness (Olbrich et al., 2009; Sander et al., 2015), and the whole-brain correlation patterns of nine brainstem and two bilateral basal forebrain regions of the ascending arousal network were compared between the two vigilance states (Edlow et al., 2024; Edlow et al., 2012; Zaborszky et al., 2008).

In addition to the vigilance-dependent FC analysis, we evaluated the ability of fMRI to reliably characterize the FC of nuclei in the brainstem and basal forebrain. Functional MRI investigations of brainstem and basal forebrain nuclei are challenging because of their small size, heterogeneity in location across individuals, and susceptibility to contamination by physiological noise due to their close proximity to major blood vessels, subarachnoid cisterns, and the ventricles (Beissner, 2015; Brooks et al., 2013). Advanced acquisition techniques, such as multi-echo sequences and 7T fMRI, may alleviate some of these limitations by improving the BOLD contrast, signal-to-noise ratio (SNR), and spatial resolution and specificity (Chang et al., 2016; Sclocco et al., 2018; Turker et al., 2021). In particular, multi-echo independent component analysis can remove non-BOLD artifacts caused by head motion and cyclic physiological noise (Kundu et al., 2013; Kundu et al., 2012). Additional preprocessing methods that estimate and regress out non-neuronal BOLD signals originating from systemic vascular effects may also improve the SNR (Brooks et al., 2013; Caballero-Gaudes and Reynolds, 2017).

We implemented a 3T multi-echo fMRI paradigm for the simultaneous EEG-fMRI dataset to mitigate SNR limitations caused by non-BOLD motion and physiological noise, and we used two large datasets of 3T and 7T single-echo fMRI from the Human Connectome Project (Smith et al., 2013) to quantify the spatial reproducibility of the static whole-brain correlation patterns of the neuromodulatory nuclei across different field strengths and acquisition methods. Because the optimal preprocessing strategy for analysis of subcortical fMRI remains an open question (Beissner, 2015; Sclocco et al., 2018; Turker et al., 2021), the FC patterns were also compared between three preprocessing pipelines designed to remove non-neuronal influences. Finally, we analyzed simultaneous fMRI and pupillometry recordings in the HCP 7T dataset to assess the reproducibility of the vigilance-dependent FC profiles of the subcortical nuclei between the EEG-fMRI and HCP 7T datasets.

## 2. RESULTS

This study included resting-state fMRI data from three datasets (see **Table 1** for a detailed description of each dataset). The first dataset consisted of simultaneous EEG and 3T multi-echo fMRI data collected at Vanderbilt University (VU 3T-ME dataset: n = 30 healthy subjects). The other two datasets consisted of 3T and 7T single-echo, multi-band fMRI from a large number of subjects in the HCP database (HCP 3T dataset: n = 375; HCP 7T dataset: n = 176) (Smith et al., 2013). Non-BOLD physiological and motion artifacts in the fMRI data were removed with multi-echo independent component analysis (ME-ICA) in the VU 3T-ME dataset (Kundu et al., 2013; Kundu et al., 2012; Turker et al., 2021) and with ICA-FIX in the HCP 3T and 7T datasets (Smith et al., 2013). fMRI signals were extracted from subcortical regions-of-interest (ROIs) involved in arousal and autonomic regulation (hereafter referred to as “arousal ROIs”). The arousal ROIs consist of monoaminergic, glutamatergic, and cholinergic nuclei in the brainstem (9 ROIs) (Edlow et al., 2024; Edlow et al., 2012) and basal forebrain (2 bilateral ROIs) (Zaborszky et al., 2008) (see **Table 2**).

**Table 1.** Demographic and technical information for the three datasets used in this study: simultaneous EEG and 3T multi-echo fMRI from Vanderbilt University (VU) and 3T and 7T single-echo fMRI from the Human Connectome Project (HCP S1200 data release).

| Dataset | VU 3T-ME | HCP 3T | HCP 7T |
| --- | --- | --- | --- |
| Number of Subjects (n) | 30 | 375 | 176 |
| Number of Sessions (n) | 45 (1-2 per subject) | 1500 (4 per subject) | 704 (4 per subject) |
| Age (mean ± SD years) | 35.1 ± 15.3 | 28.5 ± 3.8 | 29.4 ± 3.3 |
| Gender (M/F) | 14/16 | 207/168 | 70/106 |
| fMRI Modality | 3T multi-echo fMRI | 3T single-echo fMRI | 7T single-echo fMRI |
| Scalp EEG | Yes (30 subjects) | No | No |
| Respiratory belt and PPG | Yes (28 subjects) | Yes (375 subjects) | No |
| Pupillometry | No | No | Yes (145 subjects) |

**Table 2.** Seed regions-of-interest (ROIs) used for the whole-brain connectivity analysis. The ROIs are involved in arousal and autonomic regulation and consist of monoaminergic, glutamatergic, and cholinergic nuclei in the brainstem (Harvard Ascending Arousal Network [AAN] atlas Version 1.0; https://www.nmr.mgh.harvard.edu/resources/aan-atlas) (Edlow et al., 2024; Edlow et al., 2012) and basal forebrain (JuBrain Anatomy Toolbox; https://www.fz-juelich.de/en/inm/inm-7/resources/jubrain-anatomy-toolbox) (Zaborszky et al., 2008).

| Brain Area | Region-of-interest (ROI) | Main neurotransmitter systems<br>(Brown et al., 2012; Edlow et al., 2012; Scammell et al., 2017) |
| --- | --- | --- |
| Brainstem | Locus coeruleus (LC) | Norepinephrine |
| Brainstem | Dorsal raphe (DR) | Serotonin |
| Brainstem | Median raphe (MR) | Serotonin |
| Brainstem | Ventral tegmental area (VTA) | Dopamine |
| Brainstem | Periaqueductal gray (PAG) | Dopamine, GABA |
| Brainstem | Parabrachial nuclear complex (PBC) | Glutamate |
| Brainstem | Cuneiform/subcuneiform nucleus (CSC) | Glutamate, GABA |
| Brainstem | Pedunculo pontine nucleus (PPN) | Acetylcholine, glutamate, GABA |
| Brainstem | Nucleus pontine oralis (PO) | Acetylcholine, glutamate, GABA |
| Basal Forebrain | Nucleus basalis of Meynert (NBM) | Acetylcholine, glutamate, GABA |
| Basal Forebrain | Medial septum/diagonal band of Broca (MS/DBB) | Acetylcholine, glutamate, GABA |

The quality of the fMRI signals of the arousal ROIs was assessed by computing the temporal SNR (tSNR) (shown in **Supplementary Fig. 1**) from the ME-ICA denoised data in the VU 3T-ME dataset and from the ICA-FIX denoised data in the HCP datasets. The tSNR of the arousal ROIs was greater for the VU 3T-ME dataset compared to the HCP 3T and 7T datasets. In the VU 3T-ME dataset, the tSNR of the arousal ROIs was comparable to the tSNR of cortical ROIs from the Schaefer atlas (Schaefer et al., 2018). In the HCP 3T and 7T datasets, the tSNR of the arousal ROIs was lower than that of the cortical ROIs.

### 2.1. Cross-modality reproducibility of static connectivity patterns

The whole-brain static FC patterns of the arousal ROIs were estimated by computing the seed-based correlation over the entire fMRI scan duration. The seed-based correlation was calculated after removal of mean white matter (WM), deep cerebrospinal fluid (CSF), and fourth ventricle (FV) signals from the fMRI data (i.e., the mCSF/WM pipeline). The mCSF/WM pipeline is described in more detail in the **Methods** section and was performed to mitigate non-neural influences due to systemic vascular effects (Caballero-Gaudes and Reynolds, 2017; Turker et al., 2021). FC t-maps were then computed for the group average of the seed-based correlation patterns in each dataset, and the t-maps were thresholded to portray the strongest significant correlations (threshold of p_FDR_ < 0.05 and 40% of the top t-values). The static FC t-maps of the LC, cuneiform/subcuneiform nucleus (CSC), and NBM are depicted in **Fig. 1a**, and the static FC t-maps of all the arousal ROIs are provided in a Neurovault repository (available upon acceptance of this manuscript; NIFTI file format). For ease of visualization, the spatial overlap of the static FC t-maps with 16 canonical brain network templates from the FINDLAB and Melbourne atlases (Shirer et al., 2012; Tian et al., 2020) was also computed (see **Fig. 1b**).

**Fig. 1.**
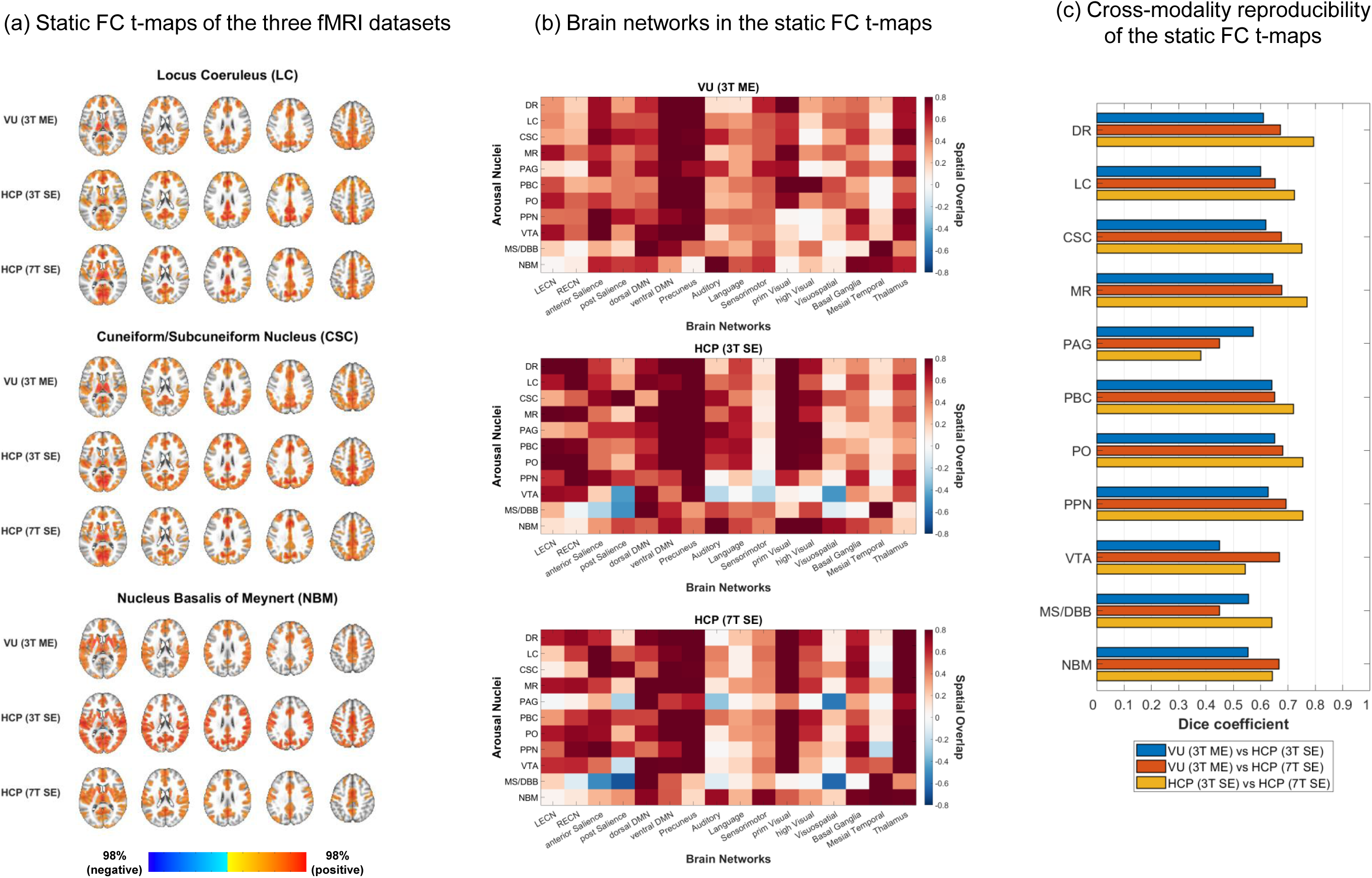
Static functional connectivity of the subcortical arousal nuclei. (a) Static functional connectivity (FC) t-maps of the locus coresuleus (LC), cuneiform/subcuneiform nucleus (CSC), and nucleus basalis of Meynert (NBM) in the VU 3T-ME, HCP 3T, and HCP 7T datasets for the mCSF/WM preprocessing pipeline. The FC t-maps were thresholded at 40% of the top t-values in the gray matter and at p < 0.05 (voxel-wise false discovery rate [FDR]-corrected over the entire gray matter volume). AFNI was used for visualization of the t-maps (@chauffeur_afni function; upper functional range set to the 98^th^ percentile). (b) Spatial overlap of the thresholded static FC t-maps of the subcortical arousal regions with 16 canonical brain network templates from the FINDLAB and Melbourne atlases (Shirer et al., 2012; Tian et al., 2020). (c) Spatial reproducibility (Dice similarity coefficient) of the thresholded static FC t-maps between the three fMRI datasets.

The Dice similarity coefficient (DSC) was used to evaluate the reproducibility of the thresholded static FC t-maps across the three fMRI modalities (see **Fig. 1c**) (Turker et al., 2021). We found that the reproducibility across all three modalities was moderate to good for all of the arousal ROIs (DSC = 0.59-0.68 [interquartile range; IQR]), except for the periaqueductal gray (PAG) between the HCP 3T and 7T datasets. The FC pattern of the ventral tegmental area (VTA) had the lowest reproducibility between the VU 3T-ME and HCP 3T datasets while the FC of the PAG and medial septum/diagonal band of Broca (MS/DBB) had the lowest reproducibility between the VU 3T-ME and HCP 7T datasets. The FC of the PAG also had the lowest reproducibility between the HCP 3T and 7T datasets.

In agreement with the moderate to good reproducibility, the thresholded FC patterns of most of the arousal ROIs were qualitatively similar between the three fMRI modalities. The LC exhibited strong positive correlations to regions of the thalamus, precuneus, basal ganglia, and salience, default mode, sensorimotor, and visual networks. The FC patterns of the other brainstem ROIs were relatively similar to that of the LC (see **Supplementary Fig. 2** for the spatial similarity of the FC patterns between the arousal ROIs). The NBM exhibited strong positive correlations to regions of the thalamus, basal ganglia, mesial temporal lobe, and salience, default mode, auditory, language, and sensorimotor networks. Notable differences between the fMRI modalities include less spatial overlap of the FC patterns of the brainstem ROIs with the sensorimotor cortex in the HCP 3T dataset and greater spatial overlap with the executive control network and higher-order visual cortex in the HCP 3T and 7T datasets.

### 2.2. EEG-based vigilance-dependent connectivity patterns

We leveraged simultaneous EEG and fMRI data in the VU 3T-ME dataset to derive vigilance-dependent FC patterns for the arousal ROIs. Time periods of alert and drowsy vigilance states were identified from the EEG data using an adapted version of the Vigilance Algorithm Leipzig (VIGALL) algorithm (see **Fig. 2a** for an illustration of the vigilance staging algorithm) (Huang et al., 2015; Jawinski et al., 2019; Sander et al., 2015). Whole-brain FC t-maps were then computed for the group average of the seed-based correlation patterns of the arousal ROIs in each state separately and for the effect of vigilance state (drowsy versus alert) on the correlation patterns. The alert, drowsy, and drowsy versus alert FC t-maps were thresholded to portray the strongest significant correlations (threshold of p_FDR_ < 0.05 and 40% of the top t-values). The vigilance-dependent FC t-maps of the LC, CSC, and NBM are depicted in **Fig. 2b**, and the vigilance-dependent FC t-maps of all the arousal ROIs are provided in the Neurovault repository. The spatial overlap of the FC t-maps with the canonical brain network templates is shown in **Fig. 2c**.

**Fig. 2.**
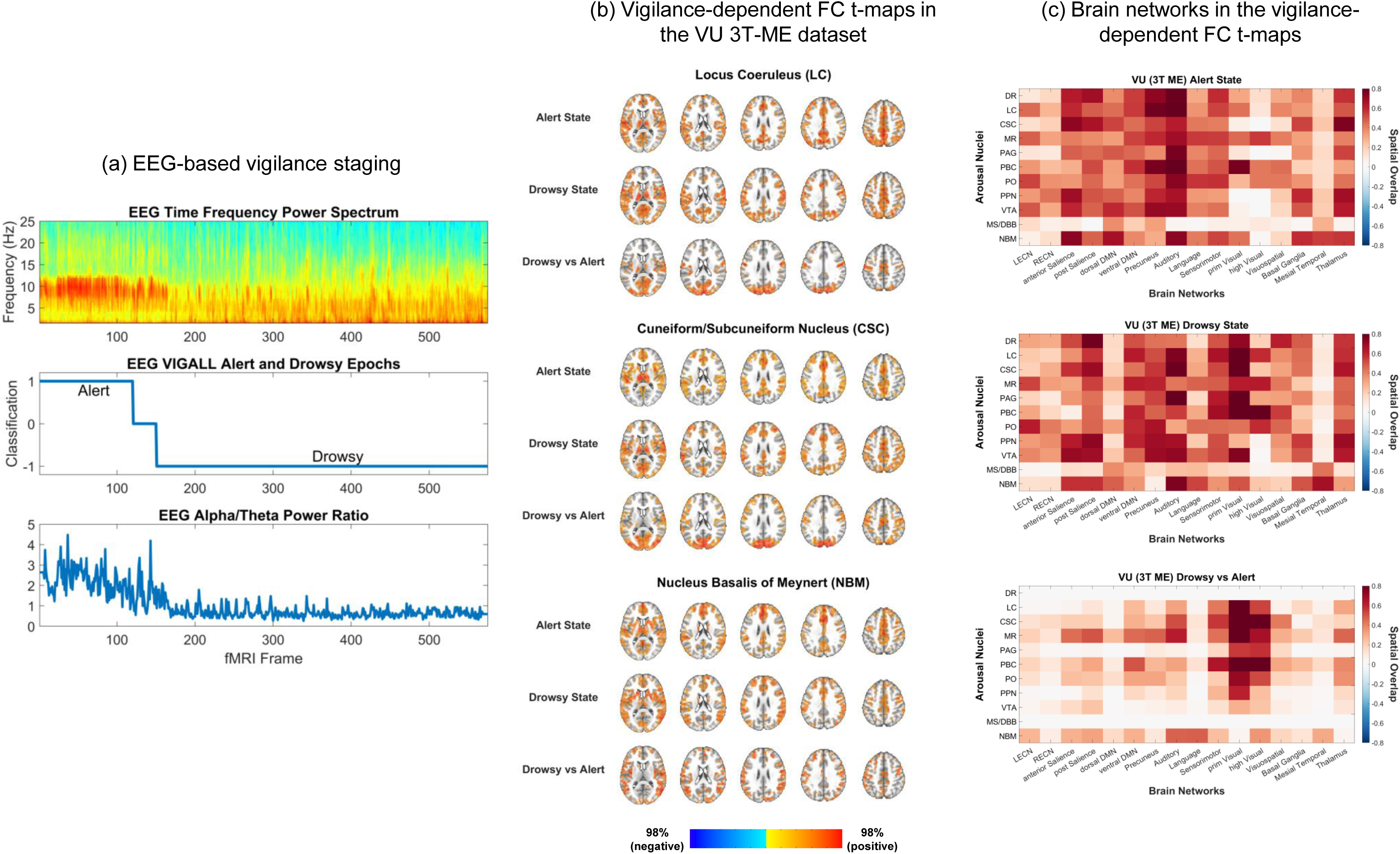
Vigilance-dependent functional connectivity of the subcortical arousal nuclei. (a) An adapted version of the Vigilance Algorithm Leipzig (VIGALL) algorithm was used to perform automatic vigilance staging of the simultaneous EEG recordings in the VU 3T-ME dataset (Huang et al., 2015; Jawinski et al., 2019; Sander et al., 2015). The accuracy of the algorithm was assessed by comparing the alert and drowsy classifications with a previously validated quantitative index of vigilance (i.e., the EEG alpha/theta power ratio) (Goodale et al., 2021; Oken et al., 2006). (b) Vigilance-dependent functional connectivity (FC) t-maps of the locus coresuleus (LC), cuneiform/subcuneiform nucleus (CSC), and nucleus basalis of Meynert (NBM) in the VU 3T-ME dataset for the mCSF/WM preprocessing pipeline. The FC t-maps were thresholded at 40% of the top t-values in the gray matter and at p < 0.05 (voxel-wise false discovery rate [FDR]-corrected over the entire gray matter volume). AFNI was used for visualization of the t-maps (@chauffeur_afni function; upper functional range set to the 98^th^ percentile). (c) Spatial overlap of the thresholded vigilance-dependent FC t-maps of the subcortical arousal regions with 16 canonical brain network templates from the FINDLAB and Melbourne atlases (Shirer et al., 2012; Tian et al., 2020).

We found that the FC of all the arousal ROIs, except for the dorsal raphe (DR) and MS/DBB, were significantly different between the alert and drowsy states. The LC, CSC, median raphe (MR), parabrachial nuclear complex (PBC), nucleus pontine oralis (PO), and NBM had the greatest vigilance-related FC alterations. In general, the arousal ROIs exhibited a stronger global correlation pattern in the drowsy compared to the alert state. The brainstem ROIs had the strongest drowsy versus alert FC differences in regions of the thalamus, precuneus, and salience, ventral default mode, sensorimotor, auditory, and visual networks while the NBM had the strongest drowsy versus alert FC differences in regions of the mesial temporal lobe and executive control, salience, ventral default mode, language, sensorimotor, auditory, and higher-order visual networks.

In the separate alert and drowsy states, the thresholded FC patterns of the arousal ROIs had similar spatial profiles as their static FC patterns. Most of the arousal ROIs had strong positive correlations to the thalamus, precuneus, and salience, default mode, auditory, and sensorimotor networks in both the alert and drowsy states. The ROIs also had strong correlations to the visual networks in the drowsy state. The FC patterns of most of the ROIs in the alert state had more spatial overlap with the dorsal default mode network than the FC patterns in the drowsy state while the FC in the drowsy state had more spatial overlap with the visual networks. The FC of most of the ROIs in both the alert and drowsy states had more spatial overlap with the auditory network compared to their static FC patterns.

### 2.3. Cross-modality reproducibility of vigilance-dependent connectivity patterns

We evaluated the cross-modality reproducibility of the state-dependent FC patterns of the arousal ROIs that had the greatest vigilance-related FC alterations in the VU 3T-ME dataset (i.e., LC, CSC, MR, PBC, PO, and NBM). An unsupervised clustering algorithm was used to derive dynamic FC states in the VU 3T-ME and HCP 7T datasets, and markers of vigilance were estimated from the simultaneous EEG data in the VU 3T-ME dataset and from the simultaneous pupillometry recordings in the HCP 7T dataset. The unsupervised clustering was performed by first computing the dynamic FC of the arousal ROIs with sliding window correlations. The k-means algorithm was then employed to spatially cluster the dynamic FC patterns of each ROI into two states (Wang et al., 2016). Whole-brain FC t-maps were derived for the group average of the sliding window correlation patterns in each state separately and for the effect of state (state 2 versus state 1) on the correlation patterns. The FC t-maps were thresholded at p_FDR_ < 0.05 and at 40% of the top t-values, and the DSC was used to evaluate the reproducibility of the single- and two-state FC t-maps between the VU 3T-ME and HCP 7T datasets. The state-dependent FC t-maps of the LC and NBM are depicted in **Fig. 3a-b**, and the FC t-maps of all the arousal ROIs (i.e., LC, CSC, MR, PBC, PO, and NBM) are provided in the Neurovault repository.

**Fig. 3.**
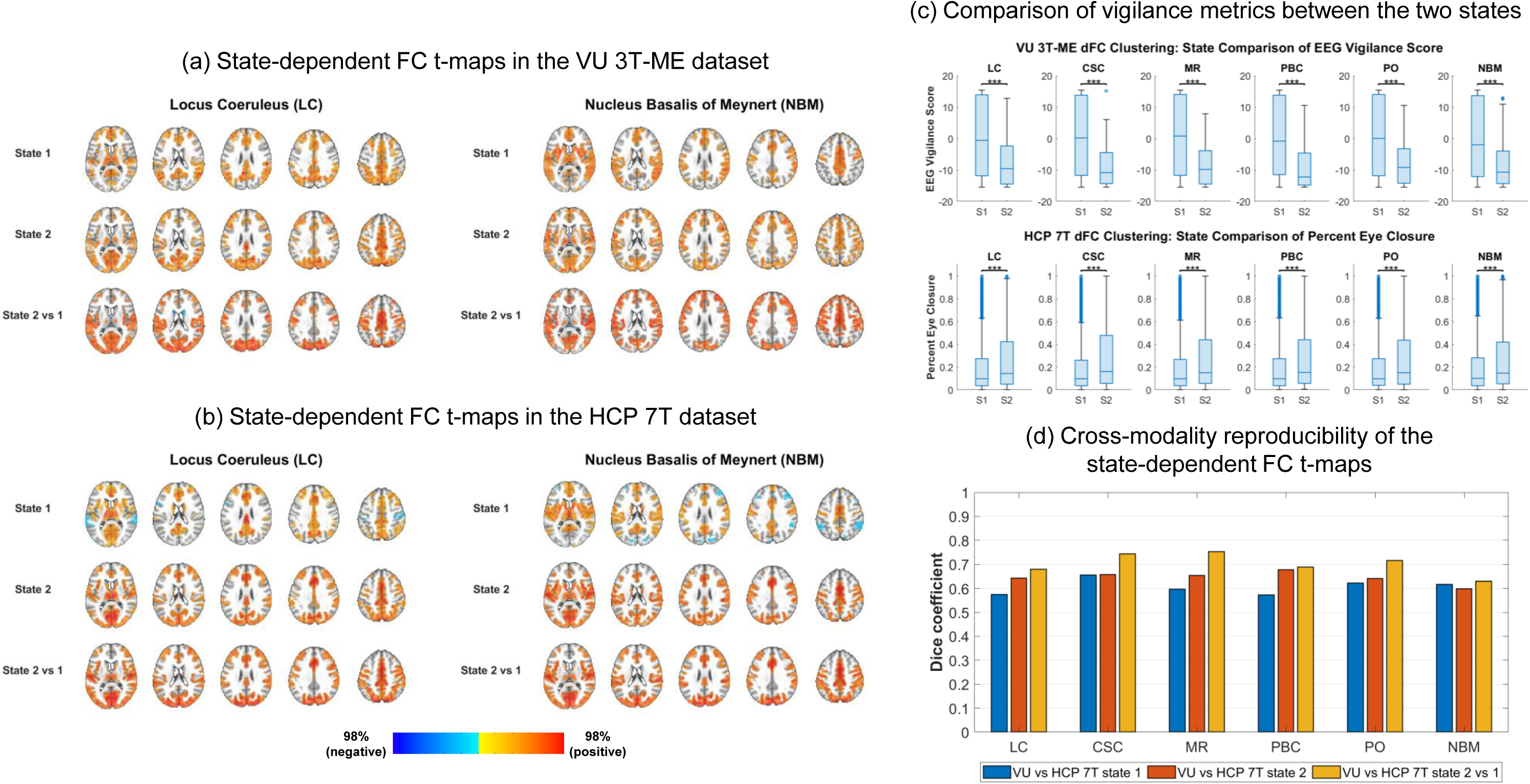
Cross-modality reproducibility of the vigilance-dependent functional connectivity. (a-b) State-dependent functional connectivity (FC) t-maps of the locus coreuleus (LC) and nucleus basalis of Meynert (NBM) in the VU 3T-ME and HCP 7T datasets for the mCSF/WM preprocessing pipeline. Unsupervised clustering of the dynamic whole-brain correlation patterns of each subcortical arousal nuclei was used to derive the two states. The FC t-maps were thresholded at 40% of the top t-values in the gray matter and at p < 0.05 (voxel-wise false discovery rate [FDR]-corrected over the entire gray matter volume). AFNI was used for visualization of the t-maps (@chauffeur_afni function; upper functional range set to the 98^th^ percentile). (c) Comparison of vigilance metrics (i.e., EEG vigilance score in the VU 3T-ME dataset and percent eye closure in the HCP 7T dataset) between the two states. Asterisks indicate a significant difference at ***p < 1e-3 (FDR-corrected across the six subcortical arousal regions). (d) Spatial reproducibility (Dice similarity coefficient) of the thresholded state-dependent FC t-maps between the VU 3T-ME and HCP 7T datasets.

In the VU 3T-ME dataset, the VIGALL-based alert/drowsy staging algorithm was used to assign a vigilance score to each time window based on the EEG data. In the HCP 7T dataset, the percent duration of eye closure was computed from the pupillometry recordings and used as a putative marker of vigilance (Abe, 2023; Shekari Soleimanloo et al., 2019; Soon et al., 2021; Wang et al., 2016). We found that, for each arousal ROI, the VIGALL score was significantly lower and the percent eye closure was significantly greater for state 2 compared to state 1 (p_FDR_ < 0.05; see **Fig. 3c**), suggesting that state 2 primarily corresponds to a state of drowsiness. The VIGALL scores of the time windows in state 1 were evenly distributed between alert and drowsy classifications (46-47% [IQR] percent alert and 47-49% [IQR] percent drowsy), suggesting that state 1 corresponds to a mixed state of alertness and drowsiness. The time windows in state 2 were primarily classified as drowsy (6-9% [IQR] percent alert and 81-87% [IQR] percent drowsy).

The single-and two-state FC t-maps had a high cross-modality reproducibility for the LC, CSC, MR, PBC, PO, and NBM (DSC = 0.62-0.68 [IQR]; see **Fig. 3d**), and the FC patterns were qualitatively similar between the VU 3T-ME and HCP 7T datasets. Similar to the EEG-derived drowsy versus alert FC patterns in the VU 3T-ME dataset, the FC of the arousal ROIs exhibited a stronger global correlation pattern in state 2 compared to state 1, with greater FC to regions of the thalamus, precuneus, and salience, ventral default mode, auditory, sensorimotor, and visual networks. Likewise, the thresholded single-state FC maps exhibited a similar correlation pattern as their EEG-derived alert and drowsy counterparts. The FC patterns in state 1 had more spatial overlap with the dorsal default mode network than the FC patterns in state 2, and the FC in state 2 had more overlap with the auditory, sensorimotor, and visual networks.

### 2.4. Influence of preprocessing on the static connectivity

In addition to the mCSF/WM pipeline, the fMRI data were preprocessed with two alternative strategies for removing systemic vascular effects (i.e., the physio and aCompCor pipelines) (Caballero-Gaudes and Reynolds, 2017). We then compared the static FC of the arousal ROIs in the VU 3T-ME, HCP 3T, and HCP 7T datasets between the three preprocessing pipelines. The aCompCor pipeline is a more aggressive method of removing signals from the WM and CSF (Behzadi et al., 2007) while the physio pipeline involves confound regression of low-frequency physiological effects associated with heart rate and respiration (Chen et al., 2020). The static FC t-maps of the LC, CSC, and NBM for the physio and aCompCor pipelines are depicted in **Supplementary Fig. 3**.

The mCSF/WM and physio pipelines resulted in largely similar static FC patterns for the arousal ROIs, and the cross-modality reproducibility of the static FC was similar for the mCSF/WM and physio pipelines (DSC = 0.59-0.68 [IQR] for the mCSF/WM pipeline and DSC = 0.56-0.62 [IQR] for the physio pipeline; see **Supplementary Fig. 4**). The aCompCor pipeline led to a global decrease in the FC strength in all three fMRI modalities, primarily in the sensory and motor networks, and resulted in significant negative correlations for most of the arousal ROIs in the HCP 3T and 7T datasets. The cross-modality reproducibility was lower for most of the ROIs in the aCompCor pipeline compared to the other pipelines (DSC = 0.44-0.60 [IQR] for the aCompCor pipeline). However, the aCompCor pipeline improved the reproducibility between the HCP 3T and 7T datasets for the PAG, MS/DBB, and NBM.

### 2.5. Influence of preprocessing on the vigilance-dependent connectivity

We also compared the EEG-based vigilance-dependent FC of the arousal ROIs in the VU 3T-ME dataset between the mCSF/WM, physio, and aCompCor pipelines. The vigilance-dependent FC t-maps of the LC, CSC, and NBM for the physio and aCompCor pipelines are depicted in **Supplementary Fig. 5**. Preprocessing the fMRI data through the physio pipeline resulted in less pronounced vigilance-related FC alterations compared to the mCSF/WM pipeline, and only the FC patterns of the CSC, MR, PBC, PO, VTA, and NBM were significantly different between the alert and drowsy states. The CSC, MR, and PBC had the greatest vigilance-related FC alterations, with similar spatial profiles as those in the mCSF/WM pipeline. Likewise, the reproducibility of the drowsy versus alert FC patterns between the mCSF/WM and physio pipelines was moderate to good for the CSC, MR, PBC, and VTA and poor for the PO and NBM (DSC = 0.37-0.61 [IQR]; see **Supplementary Fig. 6**). The reproducibility of the FC maps in the alert and drowsy states between the mCSF/WM and physio pipelines was high for all of the arousal ROIs (DSC = 0.77-0.79 [IQR]), except for the MS/DBB in the alert state.

None of the arousal ROIs had significant FC alterations between alert and drowsy states for the aCompCor pipeline, and the overall strength of the FC patterns in the alert and drowsy states was reduced compared to the mCSF/WM and physio pipelines. The reproducibility of the FC maps in the alert and drowsy states between the aCompCor and the other two pipelines was poor to moderate for most of the arousal ROIs (DSC = 0.29-0.51 [IQR]).

## 3. DISCUSSION

Using simultaneous EEG and 3T multi-echo fMRI data, we investigated the whole-brain functional network architecture of arousal regulation centers in the brainstem and basal forebrain across EEG-derived states of vigilance. Our results revealed that the FC of most of the arousal ROIs was dependent on the vigilance level, with a stronger global correlation pattern in the drowsy state compared to the alert state. These state-dependent FC patterns were replicated in an independent 7T single-echo fMRI dataset in which pupillometry was used to assess vigilance. Furthermore, we found that the vigilance-related FC alterations were reduced but not completely removed when regressing out low-frequency physiological effects modeled from respiration and heart rate signals. Finally, we demonstrated that the most dominant connections of the static FC profiles of the brainstem and basal forebrain nuclei were reproducible across 3T multi-echo, 3T single-echo, and 7T single-echo fMRI modalities.

Most of the arousal ROIs had a stronger global correlation pattern in the EEG-derived drowsy state compared to the alert state, with stronger FC to the thalamus, precuneus, and sensory and motor networks. Previous studies have shown that the amplitude of global fMRI fluctuations is increased at lower vigilance levels and is dominated by higher signal power in sensory and motor regions (Falahpour et al., 2018; Liu and Falahpour, 2020; Pourmotabbed et al., 2024; Wong et al., 2013). This fMRI signature of vigilance is conserved across multiple experimental conditions (i.e., resting-state, sleep, and sedation) (Li et al., 2023). Likewise, prior work has discovered the existence of propagating global slow waves in fMRI that are associated with arousal transitions and are more frequent in states of drowsiness and NREM sleep (Gu et al., 2021; Li et al., 2023; Liu et al., 2018; Raut et al., 2021). The vigilance-dependent FC patterns of the arousal ROIs may be influenced by the occurrence of these global slow waves, which are characterized by activation of sensory and motor cortices and co-deactivation of arousal nuclei in the thalamus, brainstem, and basal forebrain (Gu et al., 2021; Liu et al., 2018). The gamma power of intracranial EEG recordings in monkeys also exhibits a similar propagating wave topology that has been linked to cortex-wide increases in low-frequency electrophysiological activity, providing evidence for an electrophysiological basis (Gu et al., 2021; Li et al., 2023; Liu et al., 2015; Raut et al., 2021).

The vigilance-dependent FC alterations of the arousal ROIs were reduced but not completely removed when regressing out low-frequency physiological effects from the fMRI data. This indicates that changes in respiration and heart rate are associated with some but not all of the vigilance-dependent FC differences, which may be related to the role of the subcortical arousal regions in central autonomic and cardiorespiratory regulation (Benarroch, 2018; Iacovella and Hasson, 2011). Prior work has demonstrated that physiological effects in fMRI are greater at lower vigilance levels and are strongly correlated with the global fMRI signal and with fMRI signals in the thalamus, precuneus, and sensory and motor cortices (Gold et al., 2024; Ozbay et al., 2019; Yuan et al., 2013). The precuneus and sensory cortices are brain areas with a high vascular density (Bernier et al., 2018), suggesting that the physiological covariance in fMRI may partially represent systemic effects on brain vasculature (e.g., due to changes in arterial CO_2_ concentration and blood pressure) (Chen et al., 2020; Liu, 2016; Liu et al., 2017). However, studies have shown that arousal-related global activity in fMRI co-occurs with shifts in both EEG power and peripheral physiological signals (Gold et al., 2024; Gu et al., 2022; Ozbay et al., 2019). Electrophysiological oscillations in sensory and autonomic brain regions have also been observed to be coupled with cardiorespiratory activity, potentially reflecting neural interoceptive and autonomic processes (Engelen et al., 2023; Herrero et al., 2018; Kluger and Gross, 2021).

The stronger global correlation pattern of the arousal ROIs in the drowsy state suggests that neuromodulatory arousal systems may be involved in regulating global fMRI activity. These findings agree with a previous study demonstrating that inactivation of the NBM leads to suppression of global fMRI signals (Turchi et al., 2018). Neuromodulatory regulation of global fMRI activity may occur through multiple, interconnected mechanisms. Global fMRI fluctuations have been shown to be coupled to low-frequency electrophysiological oscillations and to low-frequency variations in heart rate and respiration (Gu et al., 2022; Liu et al., 2018; Ozbay et al., 2019; Pourmotabbed et al., 2024; Wong et al., 2013). These slow signal changes may be influenced by neuromodulatory control of widespread neuronal activity, brain vasculature, and autonomic function across different vigilance states. For instance, low-frequency EEG oscillations during drowsiness and NREM sleep are thought to arise due to the influence of decreased neuromodulator levels on thalamocortical activity (Brown et al., 2012; Lorincz and Adamantidis, 2017). Neuromodulator levels also mediate brain vascular tone and astrocyte activity, which can affect low-frequency electrophysiological signals via modification of interstitial ion concentrations (Ding et al., 2016; Lewis, 2021; Rasmussen et al., 2020). In addition, subcortical arousal regions are implicated in vigilance-dependent modulation of central cardiorespiratory control (Benarroch, 2018), and fluctuations in peripheral physiological activity are associated with systemic vascular effects (Chen et al., 2020; Liu, 2016; Liu et al., 2017) and entrainment of neural activity (Engelen et al., 2023; Herrero et al., 2018; Kluger and Gross, 2021).

The most dominant connections of the static FC of the arousal ROIs were reproducible across the three fMRI modalities and consisted of strong correlations to the thalamus, basal ganglia, precuneus, sensory and motor cortices, and salience and default mode networks. These brain areas partially align with the whole-brain structural connectivity profiles of the subcortical arousal nuclei. The LC has dense projections to the thalamus, sensory and motor cortices, precuneus, and salience and default mode networks (insula, cingulate gyrus, and medial prefrontal cortex) as well as sparse projections to the basal ganglia (caudate and putamen) (Zerbi et al., 2019). Studies in rodents have employed chemogenetic stimulation techniques to demonstrate that LC projections influence the FC strength of these regions (Oyarzabal et al., 2022; Zerbi et al., 2019). Moreover, our findings revealed that the static FC patterns were highly similar across the brainstem ROIs and moderately similar between the brainstem and basal forebrain ROIs. The similarity of the FC patterns may result from the reciprocal structural connections of the arousal nuclei and from reciprocal modulation of their neurotransmitter activity (Brown et al., 2012; Edlow et al., 2024).

We found that the arousal ROIs generally had strong FC to the precuneus and salience, default mode, auditory, and sensorimotor networks in both the alert and drowsy states and strong FC to the visual networks in the drowsy state. Prior studies have provided evidence for the importance of neuromodulatory arousal systems in sensory processing (Mather et al., 2016; Poe et al., 2020), which is consistent with the strong connectivity of the subcortical arousal nuclei to the salience and sensory networks. For example, the LC-norepinephrine system has been hypothesized to interact with the salience network in order to regulate selective processing of salient stimuli (Mather et al., 2016; Poe et al., 2020). Norepinephrine and LC activity have also been shown to alter visual field receptors in the occipital lobe, modulate odor detection, and enhance auditory perception (Poe et al., 2020). Furthermore, monoaminergic neuromodulators and the basal forebrain have been implicated in regulating neural activity and FC within the default mode network (Harrison et al., 2022; Kelly et al., 2009; Nair et al., 2018; Oyarzabal et al., 2022; van Wingen et al., 2014). The default mode network has been shown to be more active during states of resting wakefulness compared to externally oriented tasks (Buckner and DiNicola, 2019), and FC of the default mode network has been associated with behavioral and electrophysiological measures of drowsiness (Chang et al., 2013; Samann et al., 2011; Ward et al., 2013). In our work, the FC patterns of the arousal ROIs had greater spatial overlap with the dorsal default mode network at a higher vigilance state, indicating that interactions between the arousal nuclei and default mode network may be involved in promoting a resting wakeful state.

Unsupervised clustering of the dynamic FC of the arousal ROIs resulted in state-dependent FC patterns that were highly reproducible between the VU 3T-ME and HCP 7T datasets. The FC in the lower vigilance state was characteristic of the sensory dominated global correlation pattern observed in the EEG-derived drowsy state while the FC in the higher vigilance state exhibited strong correlations to the thalamus, precuneus, and salience and default mode networks. The global FC pattern corresponded to a lower EEG vigilance score in the VU 3T-ME dataset and to a greater percent eye closure in the HCP 7T dataset, which is consistent with the similarity of vigilance-fMRI relationships between EEG-fMRI and pupillometry-fMRI modalities (Liu and Falahpour, 2020; Soon et al., 2021). Pupil size and dilation are indices of increased arousal and have been shown to be negatively correlated with fMRI signals in sensorimotor and visual networks and positively correlated with thalamic and brainstem regions (Murphy et al., 2014; Schneider et al., 2016; Yellin et al., 2015). Spontaneous eye closures are indices of decreased arousal and have been associated with fMRI activation in the precuneus and ventral default mode, auditory, sensorimotor, and visual networks and with deactivation in the thalamus and brainstem (Ong et al., 2015; Soon et al., 2021). The fMRI activation patterns during eye closures resemble the spatial topology of global fMRI waves that occur more often at lower vigilance levels (Gu et al., 2021; Li et al., 2023; Liu et al., 2018; Raut et al., 2021). These arousal-related brain activation events may influence the dynamic FC profiles of the subcortical arousal nuclei, which have been implicated in regulating pupil activity across different vigilance states (Joshi, 2021; Larsen and Waters, 2018).

Our findings for the static FC patterns of the arousal ROIs generally agree with the results of prior studies (Bar et al., 2016; Beliveau et al., 2015; Li et al., 2014; Turker et al., 2021; Yuan et al., 2019; Zhang et al., 2016), although some discrepancies are observed primarily in the sensorimotor and visual networks. Inconsistencies between datasets may be a consequence of differences in tSNR, preprocessing strategies, sample size, and vigilance state (e.g., eyes-closed versus eyes-open and shorter versus longer scan durations). Previous work in fMRI found that the FC of the LC is only moderately concordant between multi-echo and single-echo preprocessed fMRI data (Turker et al., 2021). In our study, the tSNR of the arousal ROIs in the multi-echo fMRI dataset was greater than the tSNR in both the 3T and 7T single-echo fMRI datasets even though ICA-FIX had been applied to mitigate non-BOLD artifacts. Additionally, we found that the aCompCor pipeline reduced the overall strength of the FC maps, introduced significant negative correlations for the HCP datasets, and resulted in lower cross-modality reproducibility. Similarly, previous studies that employed aCompCor or global signal regression observed significant negative correlations in the FC patterns of the LC, DR, VTA, and NBM (Beliveau et al., 2015; Li et al., 2014; Zhang et al., 2016). Global signal regression and aCompCor aggressively remove global contributions to the correlation profiles of the arousal ROIs, and global signal regression has been shown to shift FC networks in fMRI toward negative correlations (Murphy and Fox, 2017). These negative correlations may be a byproduct of removing vigilance-related signals from the fMRI data (Liu et al., 2017; Nalci et al., 2017).

The cross-modality reproducibility of the static FC was the lowest for the VTA, PAG, and MS/DBB. The VTA had the lowest tSNR of the arousal ROIs in all three datasets, and the PAG may be more susceptible to non-neural influences because of its proximity to the cerebral aqueduct. We implemented ME-ICA in the VU 3T-ME dataset and ICA-FIX in the HCP datasets to mitigate non-BOLD cyclic physiological artifacts, and we evaluated the FC after regressing out low-frequency physiological effects modeled from WM and CSF signals (mCSF/WM and aCompCor pipelines) or heart rate and respiration signals (physio pipeline). Other studies have implemented similar preprocessing strategies (e.g., RETROICOR and removal of WM and CSF regressors) (Bar et al., 2016; Beliveau et al., 2015; Turker et al., 2021; Yuan et al., 2019). However, the optimal preprocessing pipeline remains an open question and may include novel techniques, such as masked ICA (Beissner, 2015; Maki-Marttunen and Espeseth, 2021).

Advanced methods for localization and co-registration of the arousal ROIs, such as deep learning-based segmentation and contrast enhanced structural imaging, may also improve the accuracy of the FC estimates (Doss et al., 2023; Maki-Marttunen and Espeseth, 2021; Turker et al., 2021). An important caveat is that aggressive removal of low-frequency physiological effects during preprocessing may reduce meaningful signal variance associated with arousal-related neural and neuromodulatory activity. In particular, studies have shown that low-frequency EEG oscillations are coupled to slow pulsations in global fMRI activity, CSF flow, respiration, and cardiac signals during low vigilance states, which may reflect arousal-related metabolic clearance and autonomic processes (Fultz et al., 2019; Helakari et al., 2022; Picchioni et al., 2022).

The static FC patterns of the arousal ROIs had a moderate to good reproducibility across the three fMRI modalities despite the lower tSNR of the HCP datasets compared to the VU 3T-ME dataset. The large sample size of the HCP datasets and greater number of timepoints per subject provide greater statistical power that may compensate for the lower tSNR (Helwegen et al., 2023; Smith et al., 2013). In addition, the static FC in the 3T and 7T HCP datasets tended to have higher reproducibility with each other than with the VU 3T-ME dataset, which may be attributed to several factors. Both the HCP datasets were collected in an eyes-open condition (rather than the eyes-closed condition in the VU 3T-ME dataset) and were acquired with a multi-band fMRI sequence. The HCP datasets also share some of their subject population and have a similar age range. The neocortical FC of subcortical arousal regions has been shown to be associated with age and age-related cognitive performance (Guardia et al., 2022).

Overall, the results of our study suggest that the FC of most of the arousal ROIs is influenced by dynamic variations in vigilance state. The spatial topology of the vigilance-dependent FC may reflect the role of the arousal nuclei in regulating global fMRI activity via neurobiological, autonomic, and vascular mechanisms. These findings have broad implications for studying arousal networks in healthy individuals and for clinical investigations of disrupted arousal circuits in neurological and neuropsychiatric disorders. Degeneration of cholinergic and monoaminergic neurons is a hallmark of neurodegenerative disorders such as Parkinson’s and Alzheimer’s disease (Grothe et al., 2014; Kelberman et al., 2020; Ray et al., 2018; Schmitz et al., 2016; Seidel et al., 2015), and the fMRI activity and FC of brainstem and basal forebrain nuclei have been related to cognitive outcomes in these disease groups (Mieling et al., 2023; Serra et al., 2018; Wang et al., 2023; Zeng et al., 2022). Impaired FC of subcortical arousal regions has also been observed in temporal lobe epilepsy (Englot et al., 2018; Gonzalez et al., 2023; Gonzalez et al., 2021) and traumatic brain injury (Snider et al., 2020; Spindler et al., 2021; Woodrow et al., 2023) and may contribute to excessive drowsiness, sleep disturbances, and neurocognitive deficits of attention and executive function (Englot et al., 2020; Sandsmark et al., 2017).

However, if not properly controlled for, differences in vigilance between patient and control populations can act as a confounding factor in resting-state neuroimaging studies. Likewise, modeling vigilance-related interactions in fMRI may lead to the discovery of novel neural and physiological biomarkers of arousal and neurocognitive disturbances (Bagshaw et al., 2017; Guo et al., 2023; Wang et al., 2024).

## 4. METHODS

### 4.1. Simultaneous EEG-fMRI data collection and preprocessing

This study included resting-state fMRI data from three datasets. Detailed descriptions of the datasets are provided in **Table 1**. The first dataset consisted of 20-minute sessions of 3T multi-echo fMRI collected from 36 healthy subjects (65 sessions in total) during April 1, 2021 to April 29, 2023 at Vanderbilt University (VU 3T-ME dataset). All the participants provided written informed consent, and the study protocol was approved by the Institutional Review Board of Vanderbilt University. The MRI data were acquired on a Philips 3T Elition X scanner (Philips Healthcare, Best, Netherlands) with a 32-channel head/neck coil. The BOLD fMRI data were collected in an eyes-closed resting-state condition with a 3T multi-echo, gradient-echo EPI sequence (3-mm by 3-mm in-plane ACQ resolution; 2.5-mm by 2.5-mm in-plane reconstruction resolution; 240-mm by 240-mm in-plane FOV; slice thickness = 3 mm; slice gap = 1 mm; 30 axial slices; FA = 79°; TE = 13, 31, 49 ms; TR = 2100 ms; N = 575 volumes). A high-resolution, T1-weighted structural volume was obtained for anatomical co-registration with a multi-shot, turbo-field-echo sequence (1-mm isotropic spatial resolution; 256-by-256 in-plane FOV; 150 axial slices; FA = 8°; minimum TI delay = 634.8 ms; TE = 4.6 ms; TR = 9 ms; turbo factor = 128).

Scalp EEG, respiratory, and photoplethysmography (PPG) data were recorded simultaneously with the fMRI data. MRI scanner (volume) triggers were recorded with the EEG and physiological signals for data synchronization. The scalp EEG data were collected with a 32-channel 3T MR compatible system (BrainAmps MR, Brain Products GmbH) at a sampling rate of 5 kHz, referenced to the FCz channel, and synchronized to the scanner’s 10 MHz clock. The respiratory and PPG data were collected at a 496 Hz sampling rate using a pneumatic belt and PPG transducer integrated with the MR scanner (Philips Healthcare, Best, Netherlands). The pneumatic belt was placed around the subject’s abdomen, and the PPG transducer was attached to the subject’s index finger. Data from 15 sessions were excluded due to the presence of buffer overflow errors, data transfer artifacts, or excessive noise (e.g., unremoved residual gradient artifacts) in the EEG data. Data from 5 sessions were excluded due to missing fMRI volumes and/or abbreviated scanning sessions. All the remaining 30 subjects (15 subjects with 2 sessions and 15 subjects with 1 session; 45 sessions in total) were included in the study. Out of the remaining subjects, 2 of the subjects (4 sessions) had unusable respiratory and PPG recordings and were excluded from any analyses requiring use of the physiological data.

The 3T multi-echo fMRI data were preprocessed in AFNI (https://afni.nimh.nih.gov) using the following procedure: motion co-registration with six-parameter rigid body alignment based on the middle echo (3dvolreg function), slice-timing correction (3dTshift function), and denoising with multi-echo independent component analysis (ME-ICA) (tedana 0.0.9a toolbox). ME-ICA was performed to mitigate non-neuronal artifacts in the fMRI data caused by head motion and aliased cyclic physiological noise resulting from cardiac pulsatility and respiration-induced B0-field shifts (Kundu et al., 2013; Kundu et al., 2012; Turker et al., 2021). After the ME-ICA denoising, the fMRI data were co-registered to the structural T1-weighted image and nonlinearly warped to the MNI152 template (2-mm isotropic resolution) using the Advanced Normalization Tools (ANTs) toolbox (https://github.com/ANTsX/ANTs). Additional preprocessing of the normalized fMRI data included spatial smoothing at FWHM = 3 mm (AFNI 3dFWHMx function), confound regression of potential noise signals (described in section 4.4) and Legrende polynomials up to 4^th^ order (to remove scanner drift), and bandpass filtering at 0.01-0.15 Hz.

The EEG data were denoised using the average artifact subtraction procedure of BrainVision Analyzer 2 (Brain Products, Munich, Germany) to remove MR-induced gradient and ballistocardiogram (BCG) artifacts (Goodale et al., 2021). The EEG data were then aligned to the fMRI data, down-sampled to 250 Hz, and additionally preprocessed with the EEGLAB v2020.0 toolbox of MATLAB. The additional preprocessing steps included 60 Hz notch filtering to remove powerline noise, 0.5 high-pass and 70 Hz lowpass filtering, and rejection of noisy channels (e.g., exhibiting low correlation to neighboring electrodes). The bad channel rejection was limited to at most 3 channels. After the preprocessing, the Vigilance Algorithm Leipzig (VIGALL) algorithm was implemented to stage the EEG data into five vigilance stages (described in section 4.5) (Sander et al., 2015).

The respiratory data of the subjects were contaminated with transient periods of signal dropout due to malfunction of the transducer. These periods were visually marked and replaced with NaN values to denote missing time points (3.4-6.4% [IQR] of the total scan duration). The respiratory volume (RV) time-series, matched to the fMRI sampling rate, was then computed by calculating the temporal standard deviation of the respiratory waveform in 6-s sliding windows centered at each TR (Chang et al., 2009; Chen et al., 2020). The RV at each time window was calculated using all the available time points in the window if less than 20% of the time points were missing. The RV was assigned a NaN value otherwise. For the PPG data, the peak detection algorithm of MATLAB (findpeaks function) was used to locate time points corresponding to individual heart beats, and the time-varying inter-beat-interval was computed by calculating the difference between adjacent peak times (Chang et al., 2009; Chen et al., 2020). Outliers in the inter-beat-interval time-course were identified based on a cut-off of more than 2.5 standard deviations away from the mean and linearly interpolated (0.75-1.8% [IQR] outliers out of all the time points per session). The heart rate (HR) time-series was then computed by calculating the inverse of the median inter-beat-interval in 6-s sliding windows centered at each TR.

### 4.2. Human Connectome Project (HCP) database and preprocessing

The other two datasets included in this study consisted of a subsample of healthy subjects with 3T single-echo fMRI (HCP 3T dataset) and a subsample of healthy subjects with 7T single-echo fMRI (HCP 7T dataset) from the HCP S1200 data release (Smith et al., 2013; Van Essen et al., 2012). Respiratory and PPG data were simultaneously acquired in the HCP 3T dataset, and pupillometry data were simultaneously acquired in the HCP 7T dataset. For the HCP 3T dataset, we included subjects (n = 375 subjects; 1500 sessions in total) who had four complete 14.4-minute sessions of resting-state fMRI data and whose physiological recordings were previously reported to be of good quality (Power et al., 2020; Xifra-Porxas et al., 2021). For the HCP 7T dataset, we included subjects (n = 176 subjects; 704 sessions in total) who had four complete 15-minute sessions of resting-state fMRI data.

For both HCP datasets, the fMRI data were acquired in an eyes-open resting-state condition using a simultaneous multi-slice, gradient-echo EPI pulse sequence. The imaging parameters were 2-mm isotropic spatial resolution, FA = 52°, TE = 33.1 ms, TR = 720 ms, multiband factor = 8, N = 1200 volumes for the HCP 3T dataset and 1.6-mm isotropic spatial resolution, FA = 45°, TE = 22.2 ms, TR = 1000 ms, multiband factor = 5, N = 900 volumes for the HCP 7T dataset. The data were provided after prior preprocessing had been performed with the ICA-FIX denoising pipeline (Smith et al., 2013). Briefly, the ICA-FIX pipeline included distortion and motion correction, co-registration to the subject’s structural T1-weighted image, global intensity normalization, spatial normalization to the standard MNI space, minimal high-pass filtering (cutoff = 2000 s), and ICA with the FSL tool FIX to remove non-neural spatiotemporal components (e.g., corresponding to scanner drift, head motion, and cyclic physiological noise). We additionally preprocessed the ICA-FIX cleaned data using the following procedure. The HCP 7T fMRI data were spatially down-sampled to a 2-mm isotropic resolution to match the spatial resolution of the other two datasets, and the fMRI data of both HCP datasets were spatially smoothed at FWHM = 4 mm (AFNI 3dFWHMx function), bandpass filtered at 0.01-0.15 Hz, and temporally down-sampled by a factor of 2. Confound regression of potential noise signals (described in section 4.4) was then performed on the bandpass filtered and down-sampled fMRI data.

RV and HR signals were computed from the respiratory and PPG data in the HCP 3T dataset following the same sliding window procedure described in section 4.1. The RV and HR signals were then bandpass filtered at 0.01-0.15 Hz and temporally down-sampled by a factor of 2. In the HCP 7T dataset, the pupillometry data were aligned to the fMRI data and screened for faulty recordings according to the methodology of Gonzalez-Castillo et al (Gonzalez-Castillo et al., 2022). Out of 704 sessions, 568 had available pupillometry data. Out of these 568 sessions, the pupillometry data of 20 sessions lacked TR onset information, had abbreviated recordings, or could not be loaded. Another 26 sessions had periods of eye closure greater than 90% of the recording duration, indicating potentially defective eye tracking. These 46 sessions were excluded from any analyses requiring use of the pupillometry data, leaving a total of 522 sessions (145 subjects).

### 4.3. Brain regions of interest

Seed regions-of-interest (ROIs) were defined as the 9 brainstem ROIs of the Harvard Ascending Arousal Network (AAN) atlas Version 1.0 (https://www.nmr.mgh.harvard.edu/resources/aan-atlas) (Edlow et al., 2024; Edlow et al., 2012) and the 2 bilateral basal forebrain ROIs of the JuBrain Anatomy Toolbox (https://www.fz-juelich.de/en/inm/inm-7/resources/jubrain-anatomy-toolbox) (Zaborszky et al., 2008). The brainstem ROIs consist of monoaminergic, glutamatergic, and cholinergic nuclei of the ascending reticular activating system (ARAS) involved in regulating wakefulness, alertness, and autonomic function (Scammell et al., 2017). The basal forebrain ROIs consist of cholinergic nuclei involved in cortical activation and autonomic integration (Scammell et al., 2017). A more detailed description of the seed regions is provided in **Table 2**.

For all three datasets, time-courses for the seed regions were extracted by averaging across the time-series of all the voxels in each ROI. The ROI extraction was performed on the fMRI data at the original spatial resolution in the MNI space (i.e., 2-mm for the VU 3T-ME and HCP 3T datasets and 1.6-mm for the HCP 7T dataset) without the spatial smoothing step, without the spatial down-sampling step (HCP 7T dataset), and before the confound regression pipelines (described in section 4.4). In order to evaluate the quality of the BOLD signal in each seed region, the temporal SNR (tSNR) of the seed time-courses was computed by calculating the mean of the time-course divided by the standard deviation. The standard deviation was computed for the ICA-FIX denoised signals in the HCP datasets (which includes drift removal and minimal high-pass filtering) and for the ME-ICA denoised and detrended signals in the VU 3T-ME dataset. The tSNR of the seed regions was compared to the tSNR of ROIs from the Schaefer cortical atlas (200 ROIs, 17 brain networks) (Schaefer et al., 2018) and Melbourne subcortical atlas (32 ROIs) (Tian et al., 2020). For use in the later confound regression pipelines, physiological tissue-based signals were extracted and included mean time-courses of the white matter (WM), deep cerebrospinal fluid (CSF) (i.e., first, second, and third ventricles), and fourth ventricle (FV). Masks for the gray matter (GM), WM, and CSF were obtained from the tissue-type probability maps available in FSL (https://fsl.fmrib.ox.ac.uk/fsl; 35% threshold for the GM, 50% threshold for the CSF, and 90% threshold for the WM).

### 4.4. Static functional connectivity analysis

Static FC patterns were estimated by computing the seed-based correlation of each brainstem and basal forebrain ROI to the voxels of the entire brain over the entire fMRI scan duration. The seed-based correlation was calculated after additional preprocessing was performed with three different confound regression pipelines (i.e., the mCSF/WM pipeline, aCompCor pipeline, and physio pipeline) (Caballero-Gaudes and Reynolds, 2017). The mCSF/WM pipeline involved regression of the mean WM, deep CSF, and FV signals (Turker et al., 2021); the anatomical CompCor (aCompCor) pipeline involved regression of the first five principal components of the mean WM and deep CSF signals (Behzadi et al., 2007); and the physio pipeline involved regression of low-frequency physiological effects modeled from the RV and HR signals convolved with five respiratory and five cardiac response functions (Chen et al., 2020). Before the confound regression, missing values in the convolved RV signals due to transducer malfunction were replaced with 0’s in the regression matrix.

Signals from the WM and deep CSF may contain a mixture of neuronal and non-neuronal influences (e.g., motion and systemic vascular effects) and are often removed from the fMRI data (Caballero-Gaudes and Reynolds, 2017). The FV is in close proximity to several of the brainstem nuclei and may capture non-neuronal contamination in the seed time-courses (Turker et al., 2021). Likewise, the low-frequency physiological regressors may capture non-neuronal influences due to systemic vascular effects (e.g., changes in arterial pressure and CO_2_ concentration) (Brooks et al., 2013; Chen et al., 2020). However, the physiological regressors may also covary with neuronal activity in the central nervous system involved in autonomic regulation, and regression of these signals may be detrimental to analysis of nuclei in the brainstem and basal forebrain (Chen et al., 2020). Therefore, we sought to characterize the impact of these preprocessing techniques on the FC of the seed regions. Global signal regression was not performed considering that neuromodulatory systems in the brainstem and basal forebrain may be potential neuronal contributors of global signal fluctuations in resting-state fMRI (Turchi et al., 2018; Turker et al., 2021).

For each dataset, the voxel-wise correlation values were converted to z-scores with Fisher’s r-to-z transformation, and linear mixed-effects (LME) models were fitted to the z-transformed correlation values across all the fMRI sessions using the REML method (Chen et al., 2013). The LME model per voxel was specified with the following formula:

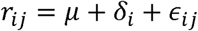

where *r_ij_* is the correlation value for subject *i* and session *j*, μ represents the group average correlation value across all subjects, δ*_i_* is a random intercept term modeling the inter-subject variance, and E*_ij_* is the residual error term modeling the intra-subject variance. We then derived t-scores for the group average correlation from the LME models. To identify brain regions with the strongest correlation to the seed ROIs, the t-maps were thresholded at 40% of the top t-values in the GM and at p < 0.05 (voxel-wise false-discovery rate [FDR]-corrected over the entire GM volume). The spatial reproducibility of the thresholded t-maps between each pair of datasets and each pair of preprocessing pipelines was evaluated using the Dice similarity coefficient (DSC) (Turker et al., 2021). The multiclass generalization of the DSC was implemented to account for positive and negative t-values in the t-maps (Taha and Hanbury, 2015). The reproducibility was scored as poor (DSC < 0.4), moderate (0.4 ≤ DSC < 0.6), and good (DSC ≥ 0.6).

For ease of visualization of the whole-brain FC patterns of the seed ROIs, we computed the spatial overlap of their thresholded FC t-maps with 16 canonical brain network templates from the FINDLAB and Melbourne atlases (Shirer et al., 2012; Tian et al., 2020). The spatial overlap values of each t-map were quantified with the Szymkiewicz-Simpson coefficient for the positive and negative t-values separately, and a signed version of the overlap coefficient was derived by taking the difference between the overlap coefficients of the positive and negative t-values.

### 4.5. EEG-based vigilance-dependent connectivity analysis

The simultaneous EEG data in the VU 3T-ME dataset provides a gold standard method of identifying time periods of alertness and drowsiness according to the spatial distribution of power changes in different frequency bands (Oken et al., 2006; Olbrich et al., 2009). For example, periods of alertness during relaxed wakefulness are characterized by dominant alpha power in the occipital region, and periods of drowsiness are characterized by greater power in the delta and theta bands (Olbrich et al., 2009). The VIGALL algorithm is an automated method for classification of scalp EEG into vigilance stages based on these spatial power distributions (Huang et al., 2015; Jawinski et al., 2019; Sander et al., 2015). In this study, the VIGALL 2.1 add-on of Brain Vision Analyzer 2 was implemented to stage each 1 second epoch of the preprocessed EEG data into five vigilance stages (i.e., A1, A2, A3, B1, B2/3) corresponding to decreasing levels of alertness. Before the vigilance staging, spherical spline interpolation was used to reconstruct EEG channels in the VIGALL standard that were not present in the data, and the EEG signals were re-referenced to the common average.

The staged EEG data were segmented into epochs of 63-s duration (30 TRs; 19 epochs per session), and a custom algorithm was used to assign each epoch into one of three vigilance states (i.e., alert, intermediate, or drowsy). First, the five VIGALL stages were converted to integer values from 1 (most drowsy) to 5 (most alert), and the Wilcoxon signed-rank test was applied to the integer values of each epoch to test for a significant difference of the median away from a (weighted) center value of 2.75. Next, a threshold of ±1.5 for the z-statistic of the signed-rank test was used to assign epochs to the three vigilance states, and adjacent epochs belonging to the same state were concatenated. The epochs were then shifted forward by 5 seconds (∼2 TRs) to account for the temporal delay between the peak BOLD response and neural activity. Our algorithm identified 21 subjects with alert epochs (n = 51 epochs; 178 ± 215 TRs per epoch) and 25 subjects with drowsy epochs (n = 75 epochs; 191 ± 208 TRs per epoch). The accuracy of the vigilance staging algorithm was assessed by comparing the alert and drowsy classifications with a previously validated quantitative index of vigilance (i.e., the EEG alpha/theta power ratio) (Goodale et al., 2021; Oken et al., 2006). The U-Sleep deep learning algorithm was also used to perform automatic sleep staging of the EEG data (Perslev et al., 2021), and we determined that the drowsy epochs primarily consisted of awake drowsy and light sleep (N1/N2) stages.

We then employed the EEG-derived vigilance states to investigate the vigilance-dependent FC of the seed regions in the fMRI data for each pipeline. The seed-based correlation of the brainstem and basal forebrain ROIs was computed for each alert and drowsy epoch, and two-state LME models were fitted to the voxel-wise correlation values across all the epochs after applying Fisher’s r-to-z transformation. The two-state LME models were specified with the following formula:

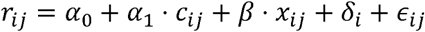

where *r_ij_* is the correlation value for subject *i* and epoch *j*, α*_0_* is the fixed intercept, α*_1_*represents the fixed effect of vigilance state *c_ij_* (i.e., alert or drowsy), and β represents a fixed slope covarying for the number of TRs per epoch *x_ij_*. We then derived t-scores for the fixed effect of vigilance state (referenced to the alert state) from the two-state LME models. For each state separately, single-state LME models were also fitted to the z-transformed correlation values:

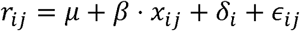

where μ represents the group average correlation value across all subjects in a single vigilance state. We derived t-scores for the group average correlation from the single-state LME models. The t-maps for the two- and single-state models were thresholded at 40% of the top t-values in the GM and at p < 0.05 (voxel-wise FDR-corrected over the entire GM volume). The DSC was then used to evaluate the spatial reproducibility of the two- and single-state t-maps between the mCSF/WM, physio, and aCompCor pipelines.

### 4.6. Pupillometry-based state-dependent connectivity analysis

The simultaneous eye-tracking recordings in the HCP 7T dataset provide a measure of vigilance and autonomic activity (Schneider et al., 2016; Wang et al., 2016). Previous studies have suggested that periods of drowsiness result in increased blink duration and more frequent and longer periods of extended eye closure (Abe, 2023; Shekari Soleimanloo et al., 2019; Soon et al., 2021). However, unlike scalp EEG, analysis of eye-tracking data does not have a clear method for automatic identification of alert and drowsy periods, and zero pupil diameter values may be confounded by instances of voluntary eye closure or device malfunction. Therefore, we characterized the state-dependent FC of the seed regions in an unsupervised manner (Wang et al., 2016), and we compared the FC patterns between the VU 3T-ME and HCP 7T datasets for the mCSF/WM pipeline.

The seed-based correlation of the brainstem and basal forebrain ROIs in the HCP 7T dataset was computed for sliding time windows of 4-minute duration and 50% overlap, and the correlation values were converted to z-scores with Fisher’s r-to-z transformation. For each ROI, the dynamic whole-brain correlation patterns were concatenated across all the 522 sessions with available pupillometry data, and k-means clustering was employed to spatially cluster the correlation patterns into different states. The distance metric was chosen to be the cityblock distance according to the recommendation of previous fMRI studies (Allen et al., 2014), and the optimal number of clusters (k = 2) was selected based on the silhouette and variance ratio criteria for a representative ROI (i.e., the LC). For the LC, the clustering analysis was repeated for window sizes of 1-minute duration. However, because no appreciable difference was observed between the clustering results for the different window sizes, 4-minute windows were selected for computational efficiency.

The percent duration of eye closure was computed for each sliding window after applying a forward shift of 4 seconds to account for the temporal delay between the peak pupil and BOLD response to neural activity (Schneider et al., 2016). The percent eye closure was calculated based on the proportion of missing (zero) pupil diameter values in each 4-minute window and includes periods of blinking and prolonged eye closure. A two-state LME model was fitted to the percent eye closure values across all the time windows to test for a significant effect of state (referenced to state 1) after applying a logit transformation to ensure normality.

The dynamic FC analysis (4-min sliding windows, 50% overlap) and k-means clustering procedure (k = 2) was repeated for each seed region in the VU 3T-ME dataset. The VIGALL-based alert/drowsy staging algorithm (described in section 4.5) was applied to the EEG data in each sliding window to derive scores of vigilance (i.e., z-scores), and a two-state LME model was fitted to test for a significant effect of state on the vigilance scores. The proportion of windows in each state that were classified as alert or drowsy was also computed after thresholding the vigilance z-scores at ±1.5.

For both the HCP 7T and VU 3T-ME datasets, LME models were fitted to the voxel-wise dynamic correlation values of each seed region to derive t-maps for the effect of state (state 2 versus 1) on the correlation values and t-maps for the group average correlation in each state separately. The t-maps were thresholded at 40% of the top t-values in the GM and at p < 0.05 (voxel-wise FDR-corrected over the entire GM volume). The DSC was then used to evaluate the spatial reproducibility of the two- and single-state t-maps between the HCP 7T and VU 3T-ME datasets.

## Acknowledgements

The analyses were conducted in part using the supercomputing resources of the Advanced Computing Center for Research and Education (ACCRE) at Vanderbilt University. Data were provided in part by the Human Connectome Project, WU-Minn Consortium (Principal Investigators: David Van Essen and Kamil Ugurbil; 1U54MH091657) funded by the 16 NIH Institutes and Centers that support the NIH Blueprint for Neuroscience Research; and by the McDonnell Center for Systems Neuroscience at Washington University.

## SUPPLEMENTARY FIGURE CAPTIONS

**Supplementary Fig 1.** Temporal signal-to-noise ratio (tSNR) of the brain regions-of-interest (ROIs) in the VU 3T-ME, HCP 3T, and HCP 7T datasets. The tSNR is averaged over all the subjects in each fMRI dataset, and the boxplots depict the distribution of the tSNR across the ROIs. The arousal ROIs include 9 brainstem regions from the Harvard Ascending Arousal Network (AAN) atlas Version 1.0 (Edlow et al., 2024; Edlow et al., 2012) and two bilateral basal forebrain regions from the Jubrain Anatomy Toolbox (Zaborszky et al., 2008). The cortical ROIs are defined from the Schaefer atlas (200 ROIs, 17 networks) (Schaefer et al., 2018), and the subcortical ROIs are defined from the Melbourne atlas (32 ROIs) (Tian et al., 2020).

**Supplementary Fig 2.** (a-b) Static functional connectivity (FC) of the subcortical arousal regions with each other in the VU 3T-ME, HCP 3T, and HCP 7T datasets for the mCSF/WM preprocessing pipeline. The FC is depicted as the Pearson correlation averaged across all the subjects in each dataset and as t-values derived for the group average correlation in each dataset. (c) Spatial similarity (Dice similarity coefficient) of the whole-brain static FC t-maps of the subcortical arousal ROIs with each other.

**Supplementary Fig. 3.** (a, c) Static functional connectivity (FC) t-maps of the locus coresuleus (LC), cuneiform/subcuneiform nucleus (CSC), and nucleus basalis of Meynert (NBM) in the VU 3T-ME, HCP 3T, and HCP 7T datasets for the physio and aCompCor preprocessing pipelines. The FC t-maps were thresholded at 40% of the top t-values in the gray matter and at p < 0.05 (voxel-wise false discovery rate [FDR]-corrected over the entire gray matter volume). AFNI was used for visualization of the t-maps (@chauffeur_afni function; upper functional range set to the 98^th^ percentile). (b, d) Spatial overlap of the thresholded static FC t-maps of the subcortical arousal regions with 16 canonical brain network templates from the FINDLAB and Melbourne atlases (Shirer et al., 2012; Tian et al., 2020).

**Supplementary Fig 4.** (a) Spatial reproducibility (Dice similarity coefficient) of the thresholded static functional connectivity (FC) t-maps between the VU 3T-ME, HCP 3T, and HCP 7T datasets for each preprocessing pipeline (mCSF/WM, physio, and aCompCor). (b) Spatial reproducibility (Dice similarity coefficient) of the thresholded static FC t-maps between the mCSF/WM, physio, and aCompCor pipelines for each fMRI dataset.

**Supplementary Fig 5.** (a, c) Vigilance-dependent functional connectivity (FC) t-maps of the locus coresuleus (LC), cuneiform/subcuneiform nucleus (CSC), and nucleus basalis of Meynert (NBM) in the VU 3T-ME dataset for the physio and aCompCor preprocessing pipelines. The FC t-maps were thresholded at 40% of the top t-values in the gray matter and at p < 0.05 (voxel-wise false discovery rate [FDR]-corrected over the entire gray matter volume). AFNI was used for visualization of the t-maps (@chauffeur_afni function; upper functional range set to the 98^th^ percentile). (b, d) Spatial overlap of the thresholded vigilance-dependent FC t-maps of the subcortical arousal regions with 16 canonical brain network templates from the FINDLAB and Melbourne atlases (Shirer et al., 2012; Tian et al., 2020).

**Supplementary Fig 6.** Spatial reproducibility (Dice similarity coefficient) of the thresholded vigilance-dependent functional connectivity (FC) t-maps between the mCSF/WM, physio, and aCompCor pipelines in the VU 3T-ME dataset.

