## Supplementary Figures for "Multimodal state-dependent connectivity analysis of arousal and autonomic centers in the brainstem and basal forebrain"

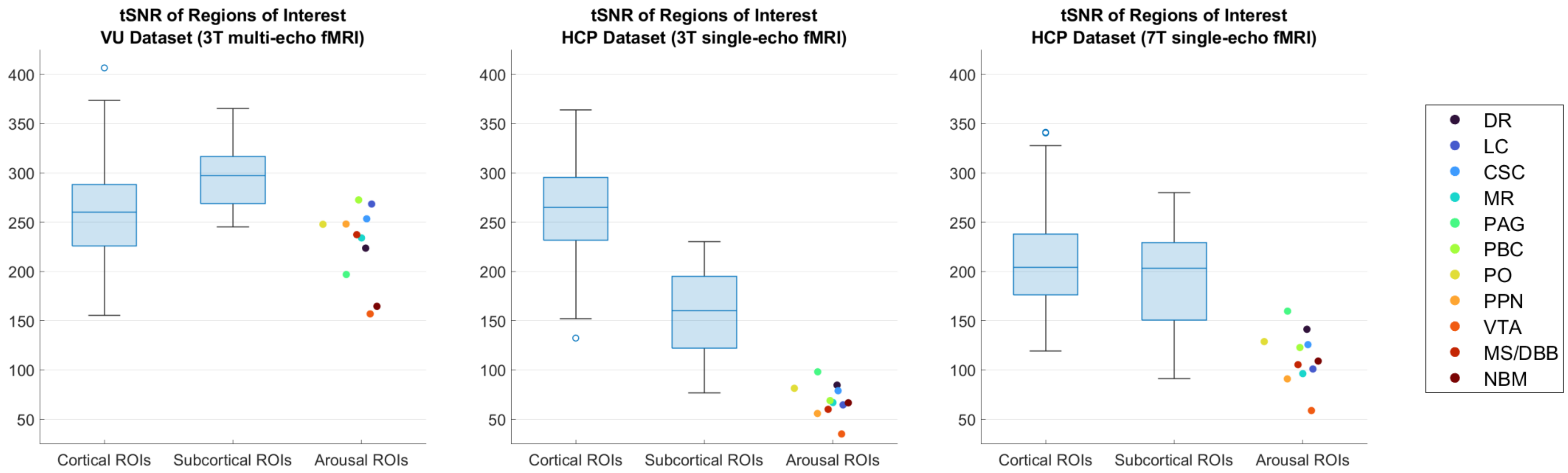


**Supplementary Fig 1.**

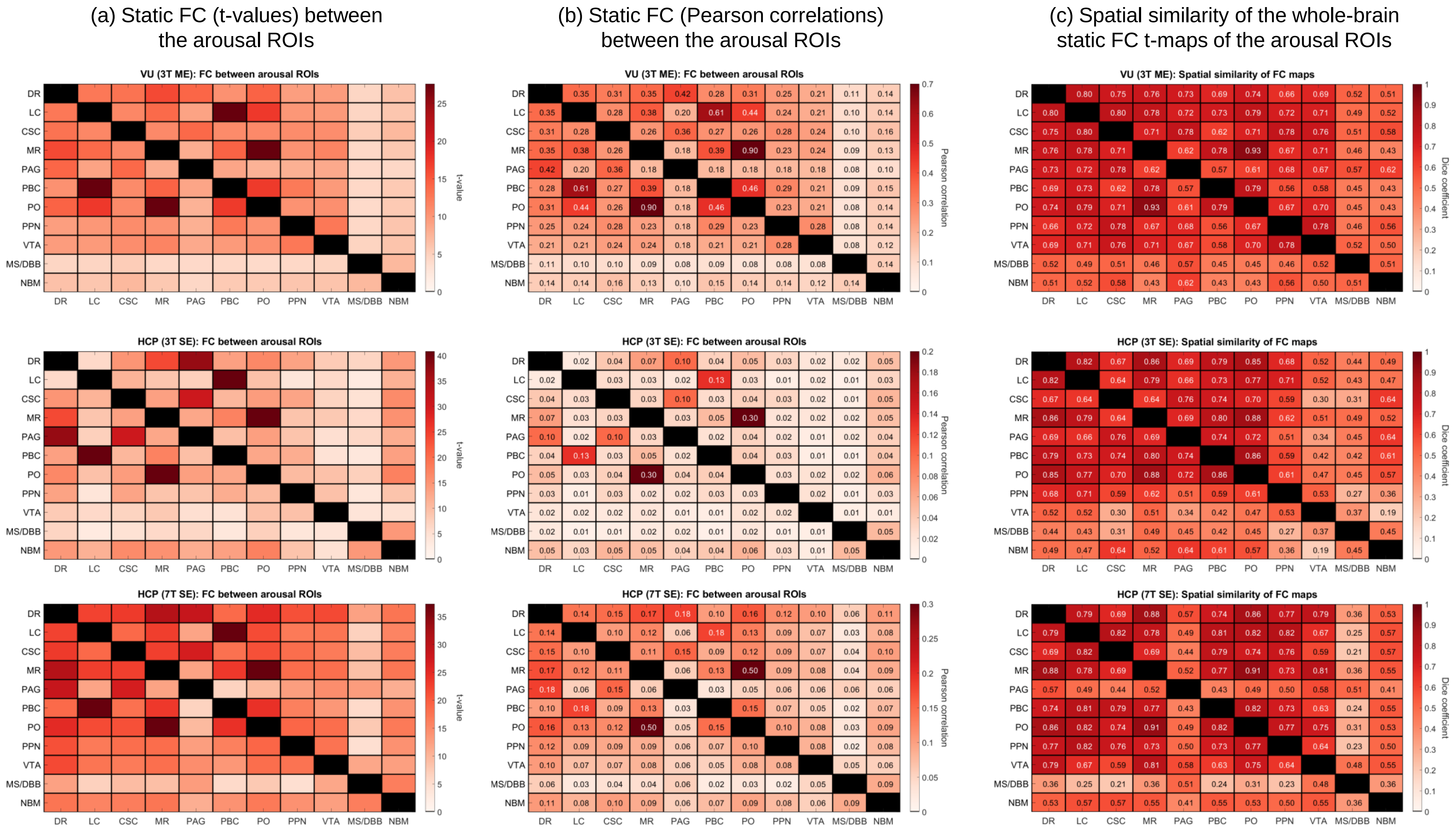


**Supplementary Fig 2.**

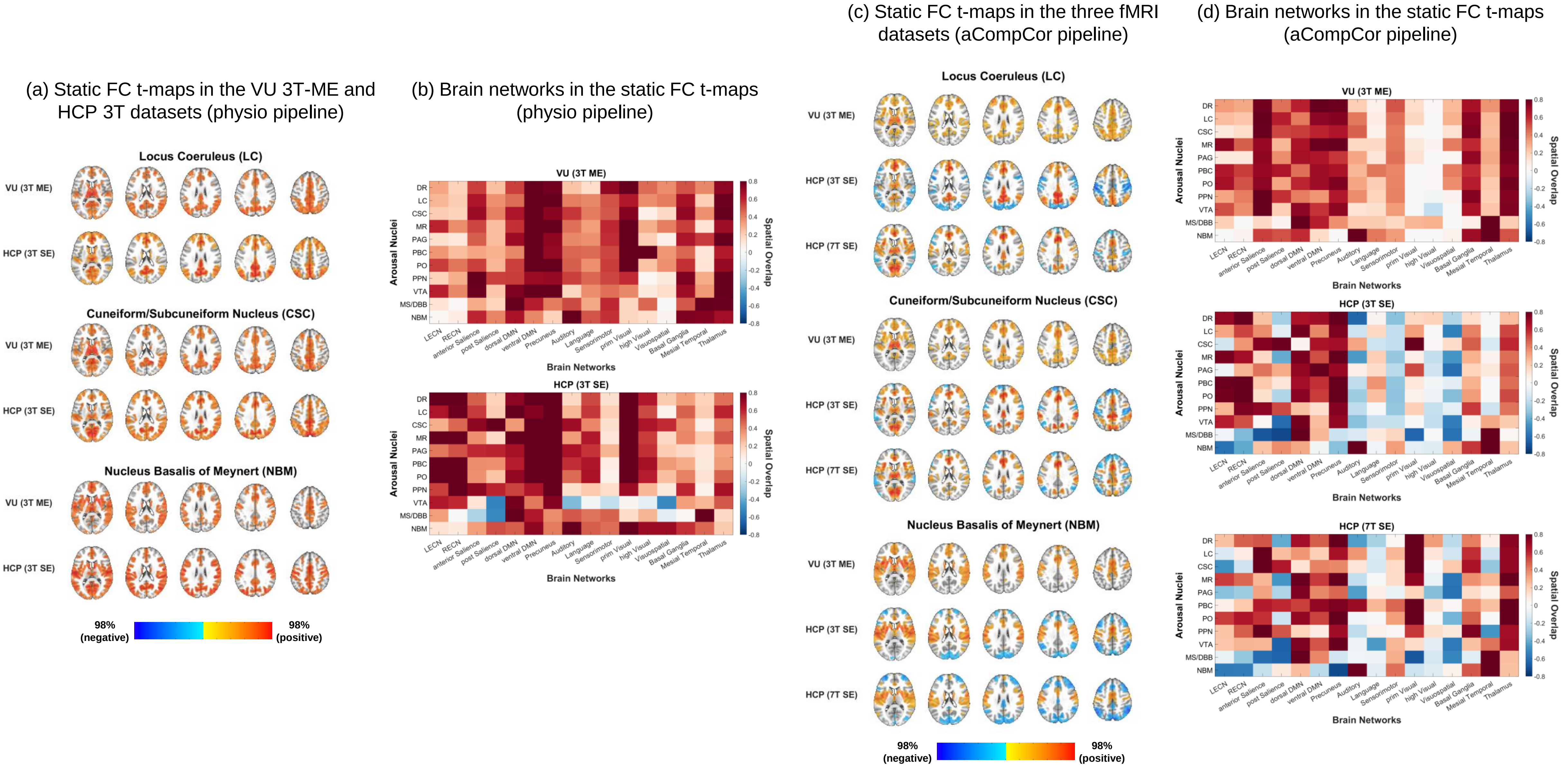


**Supplementary Fig. 3.**

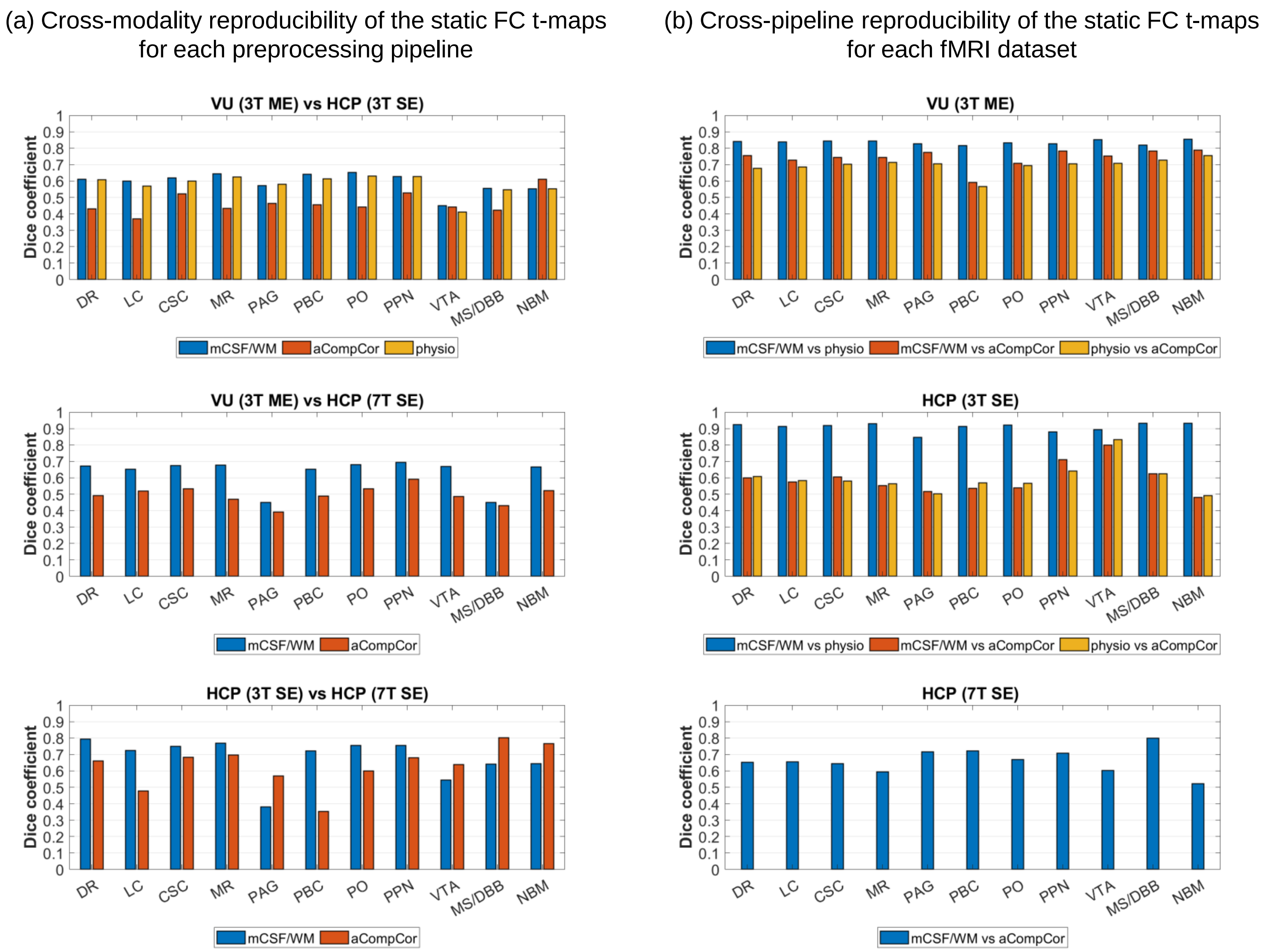


**Supplementary Fig 4.**

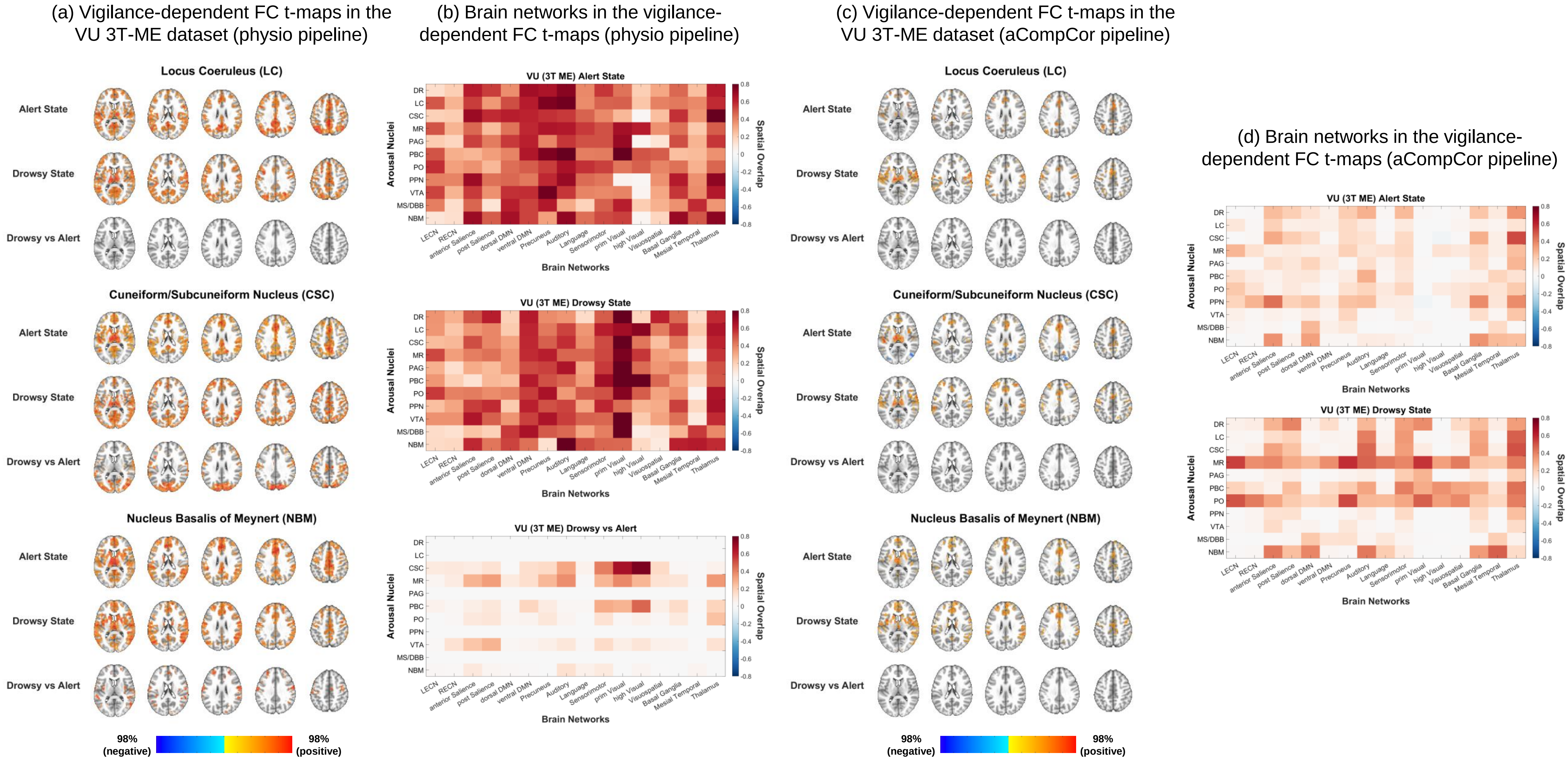


**Supplementary Fig 5.**

Cross-pipeline reproducibility of the vigilance-dependent FC t-maps in the VU 3T-ME dataset

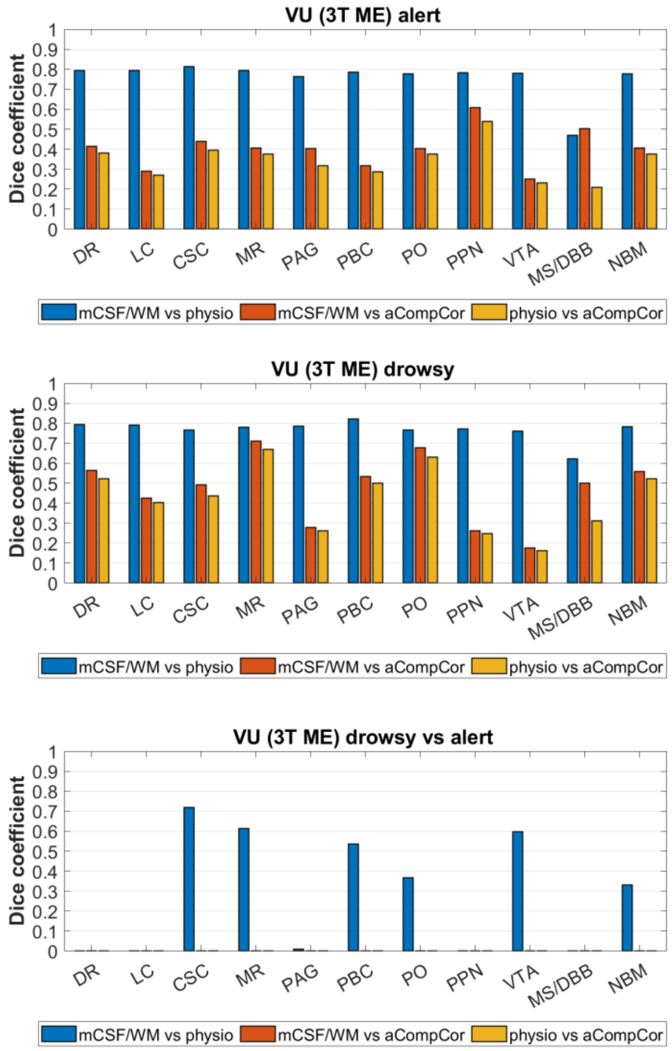


**Supplementary Fig 6.**
